# Attention to Speech Modulates Distortion Product Otoacoustic Emissions Evoked by Speech-derived Stimuli in Humans

**DOI:** 10.1101/2025.08.15.670505

**Authors:** Janna Steinebach, Tobias Reichenbach

## Abstract

Humans are remarkably skilled at understanding speech in noisy environments. While segregation of different audio streams is mostly accomplished in the auditory cortex, neural feedback connections run from the cortex to the brainstem and to the cochlea. The latter organ not only houses the mechanosensitive hair cells, but also possesses an active process enabling it to amplify sound in a frequency-dependent manner. A physiological correlate of the active process are distortion-product otoacoustic emissions (DPOAEs) that can be measured non-invasively from the ear canal. Here we employed speech-like DPOAEs that are connected to the spectral structure of voiced speech to show that these emissions are modulated by selective attention to one of two competing voices as well as by inter-modal attention. We found that speech-like DPOAEs evoked by the resolved harmonics of a voice were significantly reduced when that voice was attended as compared to when it was ignored. No such effect was observed for the unresolved harmonics of the target voice when the competing voice’s harmonics in that range were unresolved as well, indicating that attentional modulation is specific to those components of voiced speech that are spectrally resolved. Our findings support the hypothesis that the cochlea’s active process already shapes selective attention to speech in noise. Moreover, the speech-like DPOAEs that we developed open up further possibilities for investigating the contribution of the cochlear active process to auditory scene analysis in naturalistic settings.

## 1 INTRODUCTION

Understanding speech in noisy environments is an important yet highly complex human ability. In crowded settings such as restaurants or family gatherings, selective auditory attention enables listeners to focus on a single speaker while filtering out competing voices — also known as the cocktail party effect [1, 2]. This ability is essential for many aspects of social participation, but is vulnerable to hearing damage: many people with hearing impairment complain of difficulty understanding speech in background noise, even when using hearing aids [3].

The neural machinery behind selective auditory attention, including attention to speech, has been extensively studied at the level of the cerebral cortex [4, 5, 6, 7, 8]. However, anatomical and physiological evidence points to substantial descending feedback from the auditory cortex to the auditory brainstem and to the cochlea [9, 10, 11]. Through these neural feedback loops, subcortical processing centers may contribute to accomplishing the cocktail party effect. Several studies have indeed found that neural responses from the brainstem, in particular frequency-following responses, can be modulated by selective auditory attention, although others did not find an attentional effect [12, 13, 14, 15, 16, 17].

The cochlea – the sensory organ of hearing in which sound vibrations are converted into electrical signals – may already contribute to selective auditory attention as well. This fascinating organ spatially decomposes a complex sound such as speech into its individual frequency components, following a tonotopic map in which high frequencies are detected near the organ’s base, and lower frequencies progressively further towards the apex [18, 19].

In addition, the cochlea possesses an active process through which it amplifies weak sounds [20, 21]. This mechanical amplification is provided by outer hair cells and can be reduced through activation of the medial olivocochlear (MOC) fibers – efferent connections that innervate the outer hair cells [22, 23]. Because each MOC fiber is tuned to a narrow frequency band, and because the innervation of the cochlea by these fibers displays a tonotopic arrangement, the gain of the active process can potentially be regulated by the brain in a frequency-dependent manner.

The active process is accompanied by a nonlinear response that gives rise to otoacoustic emissions (OAEs). These can be recorded from the ear canal and serve as a non-invasive measure of the amplification gain. OAEs have indeed been used to assess the contribution of the cochlea to selective attention, although with inconclusive results [24, 25, 26, 27, 28, 29, 30]. A limitation of these studies was that they either did not involve naturalistic sounds such as speech that facilitate attention, or that they elicited OAEs in a manner that was not directly related to the auditory signal that the participants were asked to attend.

Computational models have shown that selective attention to a speech signal in noise may be supported by frequencyspecific modulation of the cochlear active process [31, 32]. Most parts of speech are voiced, with the energy carried by the fundamental frequency and its many higher harmonics. The lower harmonics, up to the 10th, can be spatially resolved in the cochlea, that is, they cause peaks at significantly distinct locations [33, 34, 35, 36].

A compelling hypothesis is that the cochlear amplifier selectively enhances the resolved harmonics of a target speech and suppresses the spectral bands that lie inbetween. This mechanisms would thus reduce background noise already at the level of cochlear activity. Because unresolved harmonics cannot be spatially differentiated in the cochlea, this mechanism should not be able to operate for these.

Here, we set out to test this hypothesis. We employed distortion product otoacoustic emissions that were evoked by certain higher harmonics of the voiced parts of speech (speech-like DPOAEs). As the fundamental frequency of natural speech varies over time, the stimuli used to generate speech-like DPOAEs, as well as the DPOAEs themselves, were not pure tones, but instead had instantaneous frequencies that varied over time in proportion to the fundamental frequency of the source signal. The amplitude of the stimuli varied as well, in particular, it was zero during voiceless parts of the speech signal or during silences. We recently developed and corroborated this approach [37].

We elicited and recorded speech-like DPOAEs from one ear while two competing talkers were presented to the contralateral ear. Subjects were instructed to attend either one of the two talkers or to read a text in front of them (visual attention). We then evaluated how the speech-like DPOAEs, in particular those related to resolved and unresolved harmonics, were affected by the attentional focus.

## 2 MATERIALS AND METHODS

### 2.1 Experimental design

We utilized a single- and a competing-speaker paradigm. In the single-speaker recordings, an audiobook either spoken by a female or by a male voice was presented to the right ear of the participant. Subjects were asked to focus their attention on the single voice.

In the competing-speaker scenario, two audiobooks, one spoken by a women and the other by a man, were added together and presented to each subject’s right ear (Fig. 1). Subjects were then instructed to attend either the female or the male voice. In the following, we refer to these three attentional conditions as attended female voice (Att. F) and attended male voice (Att. M). To test effects of intermodal attention, a visual distractor was introduced as well and attended upon prompt; participants then read a story displayed in segments on a screen while ignoring both audio streams. This attentional condition will be referred to as attended visual distractor (Att. V).

**Figure 1.**
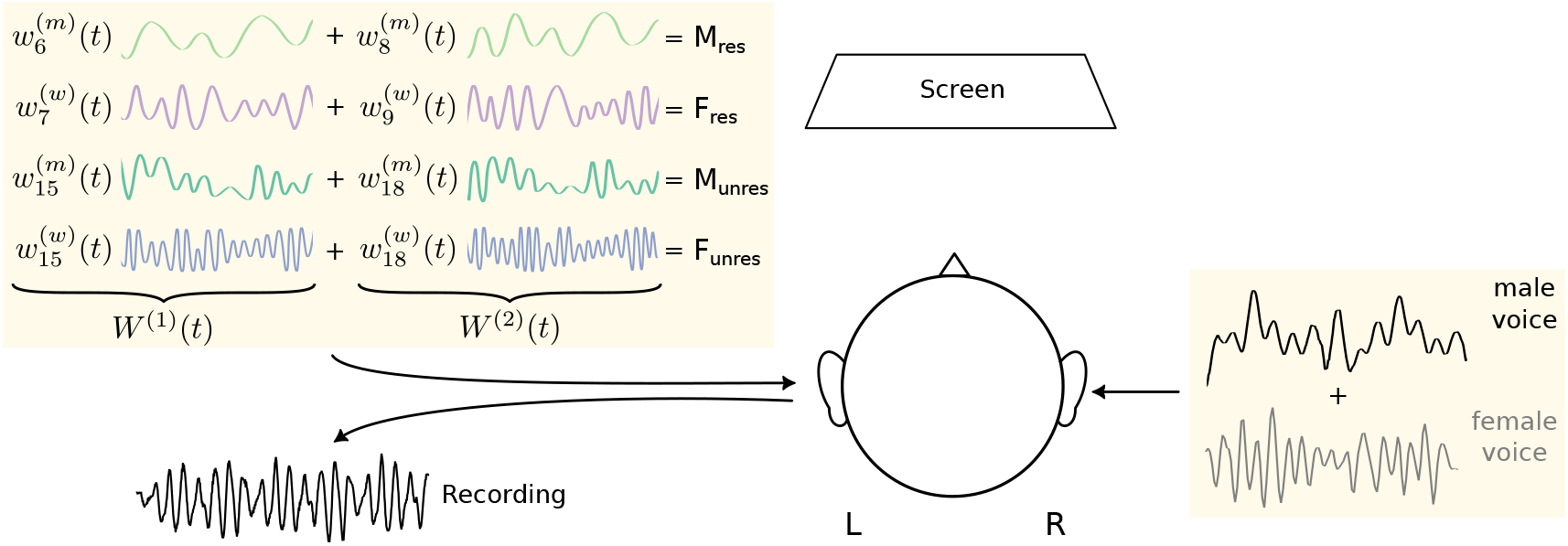
Experimental setup. Two audiobooks, one spoken by a women with fundamental frequency 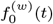 and one narrated by a man with fundamental frequency 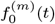, were simultaneously presented to the right ear. Speech-like DPOAEs were recorded from the left ear. They were elicited by four pairs of waveforms. The first stimulus pair, *M*_res_, consisted of two waveforms 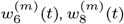 based on resolved harmonics of the male voice (frequencies 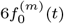 and 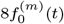. The second stimulus pair, *F*_res_, comprised two waveforms 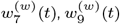 following resolved harmonics of the female voice (frequencies 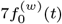 and 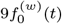. The third and fourth stimulus pair, *M*_unres_ and *F*_unres_, were waveforms 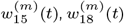 resp. 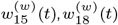 derived from unresolved harmonics of the male resp. female voice (frequencies 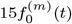 and 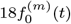 resp. 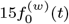 and 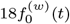. Participants directed their attention either to one of the two talkers or visually to a continuously presented story on a screen.

Speech-like DPOAEs were elicited by waveforms derived from the speech signals. To enable clean simultaneous measurement of four different speech-like DPOAEs, these were evoked and measured from the contralateral, i.e. the left, ear.

The presentation of the audiobooks together with the recordings of the speech-like DPOAEs were segmented into two-minute trials, each followed by three comprehension questions and a rating of perceived mental effort to ensure task engagement. For the auditory conditions (attending the female resp. male voice), mental effort was equivalent to the listening effort. In order to include assessment of the effort for the third, i.e. the visual, condition, the term mental effort was chosen.

### 2.2 Participants

Speech-like DPOAE measurements were conducted with *N* = 40 participants (21 female, 19 male), aged 18–31 years (mean age *±* SD: 25 *±* 3 years). Inclusion criteria were right-handedness, native German proficiency, and the absence of neurological or hearing impairments. One subject was excluded since their comprehension score was below chance level, and another due to a faulty microphone recording. 38 participants were thus included in the final analysis.

All procedures were approved by the ethics board of the University Hospital Erlangen (registration 133-12B) and were conducted in accordance with institutional regulations. Informed consent was obtained from all participants.

### 2.3 Speech signals

Two audiobooks were synthesized with a text-to-speech engine (ElevenLab, U.S.A.) using the voice “Matilda” for the female speaker and the voice “Brian” for the male one [38]. We chose voice parameters that yielded a large separation between the fundamental frequencies of the two voices, which resulted in an average fundamental frequency of 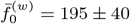 Hz for the female voice and 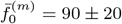 Hz for the male voice (mean *±* SD).

The texts for the audiobooks were taken from “Eine Frau erlebt die Polarnacht” [39] (Book A) and from “Darum” [40] (Book B). Each text was synthesized both with the female and the male voice.

The amplitudes of the speech signals were normalized and scaled such that the root-mean-square amplitude was (2.0 *±* 0.1) *×* 10^−3^ for both the male- and female-spoken texts. Using PsychoPy [41], the presentation level of the speech stimuli was adjusted such that it reached an average sound pressure level of 37 db SPL over the two minute segments.

### 2.4 Visual distractor

The visual distractor consisted of text excerpts from a third book, “Frau Ella” [42] (Book C). The text was rendered as a video in which short paragraphs appeared word by word at a comfortable reading pace and were displayed centrally on a computer screen.

### 2.5 Stimuli for eliciting speech-like DPOAEs

Pure-tone DPOAEs were elicited by two primary frequencies, *f*_1_ and *f*_2_. The lower-sideband cubic distortion product 2*f*_1_ − *f*_2_ is the strongest, and is maximal at a ratio of *f*_2_*/f*_1_ ≈ 1.2. For its measurement we employed *f*_1_ = 1 kHz and *f*_2_ = 1.2 kHz.

The stimuli to elicit speech-like DPOAEs were computed using an approach that we developed recently [37].

For the voiced segments of each speech signal, we first computed the fundamental waveform *w*_0_(*t*), which follows the time-varying fundamental frequency *f*_0_(*t*) of the source signal. This was achieved by applying a zero-phase, sixth-order IIR bandpass filter centered on the mean fundamental frequency 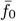, with corner frequencies of *±*0.5 standard deviations around 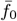. The mean fundamental frequency was estimated using the probabilistic YIN algorithm implemented in the librosa library [43]. The fundamental waveform *w*_0_(*t*) was normalized by z-scoring.

Based on *w*_0_(*t*), waveforms for the harmonic overtones *n* and *m* (*n* < *m*) were constructed such that their instantaneous frequencies equaled *nf*_0_(*t*) and *mf*_0_(*t*), respectively. To do so, we computed the analytic representation of the fundamental waveform using the Hilbert transform *H*[*w*_0_(*t*)]:

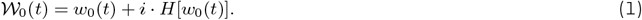

The fundamental waveform can then be expressed as the real part of the complex signal:

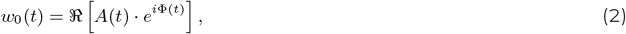

in which *A*(*t*) = |*W*_0_(*t*)| denotes the signal amplitude, and Φ(*t*) = arg[*W*_0_(*t*)] specifies its instantaneous phase. Harmonic waveforms *w*_*n*_(*t*) and *w*_*m*_(*t*) were obtained by multiplying the phase Φ(*t*) by the desired harmonic number and taking the real part:

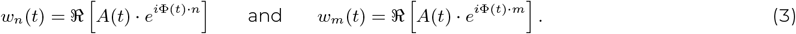

The instantaneous frequencies of the two elicitor waveforms are thus *nf*_0_(*t*) and *mf*_0_(*t*). Consequently, the lower-sideband cubic distortion product they generate exhibits an instantaneous frequency of (2*n* − *m*)*f*_0_(*t*), corresponding to the waveform *w*_2*n*−*m*_(*t*). The resulting speech-like DPOAE was identified by cross-correlating *w*_2*n*−*m*_(*t*) with the microphone recording.

To assess attentional modulation of cochlear activity at both resolved and unresolved harmonics, we designed four pairs of stimulus waveforms: (1) the stimulus *F*_res_: two waveforms 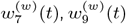 corresponding to the resolved harmonics of the female voice, (2) the stimulus *F*_unres_: two waveforms 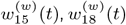 corresponding to the unresolved harmonics of the female voice, (3) the stimulus *M*_res_: two waveforms 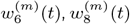 corresponding to the resolved harmonics of the male voice, and (4) the stimulus *M*_unres_: two waveforms 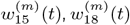 corresponding to the unresolved harmonics of the male voice (Table 1).

**Table 1.** The harmonic numbers *n* and *m* for the different stimuli, with the harmonic index 2*n* − *m* of the resulting lower-sideband cubic distortion product and the ratio *m/n*.

| | $n$ | $m$ | $2n - m$ | $m/n$ |
| --- | --- | --- | --- | --- |
| $F_{\text{res}}$ | 7 | 9 | 5 | 1.29 |
| $F_{\text{unres}}$ | 15 | 18 | 12 | 1.2 |
| $M_{\text{res}}$ | 6 | 8 | 4 | 1.33 |
| $M_{\text{unres}}$ | 15 | 18 | 12 | 1.2 |

We employed different harmonics *n* and *m* for the resolved harmonics of the female and the male voice in order to avoid unwanted correlations between the stimulus waveforms and the DPOAE waveforms. Moreover, we tried to achieve a ratio of *m/n* ≈ 1.2 for optimal distortion product generation. For unresolved harmonics, the frequency spacing was sufficiently large to allow the same harmonic pairs to be used for male- and female-related stimuli without inducing unwanted correlations between the speech-like DPOAE and stimulus waveforms.

For the simultaneous presentation of all four harmonic pairs, all waveforms corresponding to the lower harmonic number *n* were summed to form the waveform *W* ^(1)^(*t*), and all waveforms corresponding to the higher harmonic number *m* were summed separately to form the waveform *W* ^(2)^(*t*). These two waveforms were delivered to the ear canal via two independent loudspeakers.

### 2.6 Experimental Setup

Experiments were conducted in a sound-proof, semi-anechoic chamber. Stimulus presentation and data acquisition were automated through PsychoPy [41]. Instructions were displayed on a screen; responses were given via mouse click.

The sound stimuli were presented at 44.1 kHz using a high-performance sound card (RME Fireface 802) and delivered through an extended-bandwidth otoacoustic measurement system (ER10X, Etymotics, U.S.A.), equipped with one microphone and three speakers per ear. Custom ear tips ensured optimal probe fit. Audiobooks were presented to the right ear while stimuli presentation and speech-like DPOAE recordings were conducted in the left ear. For each stimulus, the two waveforms *W* ^(1)^(*t*) and *W* ^(2)^(*t*), each consisting of the sum of the four harmonic waveforms corresponding to the lower and higher harmonic numbers *n* and *m* respectively, were played through different speakers to avoid hardware-induced distortion.

Stimuli were delivered directly into the ear canal. The presentation level was adjusted so that, when averaged across each trial of approximately two minutes, the resulting mean sound pressure level in the ear canal was 37 dB SPL. All participants reported this level as comfortable. It was intentionally kept low to avoid eliciting the middle-ear muscle reflex [44, 45].

### 2.7 Experimental Routine

Each story segment lasted about two minutes. After each such two-minute trial, participants answered three comprehension questions and rated the perceived mental effort on a 13-point Likert scale.

The experiment began with a two-minute pure-tone DPOAE measurement to verify DPOAE detectability. DPOAEs could be recorded in all participants.

Next, speech-like DPOAEs were recorded in a single-speaker scenario, using either the male and the female voice in isolation. Speech-like DPOAEs to both the resolved and the unresolved harmonics of the corresponding voice were measured simultaneously to confirm that both speech-like DPOAEs could be measured concurrently.

The main part of the experiment consisted of a competing-speaker scenario in which both the female and the male voice were presented simultaneously. Participants were instructed to direct their attention either to the female speaker (Att. F), to the male speaker (Att. M), or to the visual task (Att. V). The target audio thereby always contained the story of Book A, spoken either by the male or the female voice, allowing participants to follow a continuing story when switching attention between speakers. In the Att. V. condition, participants ignored both audio streams and focused on reading Book C, presented word-by-word on a monitor. During the two auditory-attention conditions (Att. F and Att. M), the text from Book C was also shown on the monitor; however, participants were allowed to choose whether to look at it or not, depending on whether they found it distracting from the auditory task. During each of the conditions, the waveforms *W* ^(1)^(*t*) and *W* ^(2)^(*t*) comprising the four stimulus pairs *F*_res_, *F*_unres_, *M*_res_ and *M*_unres_ were presented to elicit the four respective speech-like DPOAEs.

To verify that the measurement equipment did not contribute to the speech-like DPOAEs, out-of-ear control measurements were conducted. The probe was placed outside the ear canal in the center of the recording room, with all reflective surfaces avoided to minimize acoustic feedback. No speech-like DPOAEs emerged in that case.

### 2.8 Analysis of speech-like DPOAEs

Hardware-induced delays were estimated per trial by cross-correlating the stimulus waveforms with the microphone recording. The recordings were corrected for the delays before further analysis.

Speech-like DPOAEs were computed by cross-correlating each of the four speech-like DPOAE waveforms *w*_2*n*−*m*_(*t*) with the microphone recording. To compensate for potential phase shifts between the otoacoustic emission and the waveform *w*_2*n*−*m*_(*t*), we computed the complex cross-correlation. Its real part corresponds to the correlation between the real component of the analytic representation of *w*_2*n*−*m*_(*t*) and the microphone signal, whereas the imaginary part corresponds to the correlation with the imaginary component of the analytic representation. The envelope of the complex cross-correlation was then obtained as the absolute value of the resulting complex-valued correlation.

Grand averages were computed by averaging the envelopes of the cross-correlations across all two-minute trials, distinguishing between the three attentional conditions and the four stimulus types. A peak in the grand average was considered significant if it exceeded the maximum of the noise. The noise level was thereby determined from the values of the envelope of the correlation coefficients at time lags of −750 to −70 ms and from 70 to 750 ms, that is, at delays at which no speech-like DPOAEs should occur.

To detect speech-like DPOAEs at the level of individual two-minute trials, the expected window for peak delays was defined as 1 *±* 3 ms (mean *±* SD), based on the grand averages of the stimuli *F*_*res*_ and *F*_*unres*_. Stimuli corresponding to the male voice were excluded for the determination of this window: the grand average for the *M*_*res*_ stimulus did not yield reliable results, and we wanted to prevent an imbalance between resolved and unresolved harmonics, such that we disregarded the stimulus *M*_*res*_ as well. A peak from an individual trial within this window was considered signifi-cant if it exceeded the 97th percentile of the noise. The percentage of significant trials was then computed for each stimulus type.

To compare speech-like DPOAEs across attentional conditions, the cross-correlation coefficients at the grand average peak delay were extracted for each two-minute trial, matched per stimulus type and attentional condition, and averaged per participant. Single-speaker comparisons used unpaired *t*-or Mann–Whitney *U* tests. For the data obtained from the competing-speaker scenario, paired *t*-tests or Wilcoxon signed-rank tests were used, with outlier exclusion when justified.

To evaluate differences between speech-like DPOAEs evoked by resolved and unresolved harmonics, peak delays per individual two-minute trials were extracted for all significant peaks, matched per stimulus type and condition and averaged per participant. Statistical comparisons used unpaired *t*-or Mann–Whitney *U* tests.

All *p*-values were corrected for multiple comparisons using the False Discovery Rate correction.

## 3 RESULTS

### 3.1 Comprehension scores and mental effort

Speech comprehension was quantified as the percentage of correct answers, and mental effort was rated using mean values on a Likert scale from 1 to 13 (low to high). Values are given as mean *±* SD. Statistical significance was assessed using Wilcoxon signed-rank tests.

In the single-speaker measurements, comprehension scores were high: 96.5 *±* 0.1% for the female voice and 95.6 *±* 0.1% for the male voice, with no significant difference between them.

The comprehension scores were slightly lower in the competing-speaker condition: 94.5 *±* 0.1% when attending the female voice, 91.1 *±* 0.1% for the male voice, and 93.9 *±* 0.1% when reading the text. Again, no significant differences were found, suggesting that all conditions were similarly comprehensible.

Regarding mental effort, the female and male voices were perceived as similarly demanding in the single-speaker condition, with ratings of 4.0 *±* 2.3 versus 4.4 *±* 2.4 and no significant difference.

In the competing-speaker condition, perceived effort varied significantly depending on the attentional focus. Attending the visual distractor was rated easiest at 5.7 *±* 2.2; lower than when attending the male voice (*p* < 0.001) and lower than when attending the female voice (*p* < 0.05). In contrast, attending the male voice was rated hardest at 7.3 *±* 1.7; with *p* < 0.001 when compared to other conditions. Attending the female voice fell inbetween at a value of 6.5 *±* 1.8.

### 3.2 Measurement of speech-like DPOAEs

To measure a particular speech-like DPOAE, we computed a waveform *w*_2*n*−*m*_(*t*) that corresponded to the lower-sideband cubic distortion product of the pair of harmonics that was used for stimulation. As an example, for the stimulus *F*_res_ we utilized waveforms 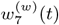 and 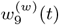, yielding 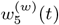 as the lower sideband cubic distortion product.

The speech-like DPOAE waveform *w*_2*n*−*m*_ was then cross-correlated with the microphone recording (Fig. 2A-C). We computed the envelope of the cross-correlation to obtain, in most subjects, a single peak at a delay between 0 − 3 ms. A peak can be interpreted as a successful measurement of a speech-like DPOAE, with a delay that corresponded to that of the peak. Examples of individual trials with large and moderate peak amplitudes, as well as a trial without a significant peak, are shown in Fig. 2A-C, illustrating the variability in speech-like DPOAE morphology.

**Figure 2.**
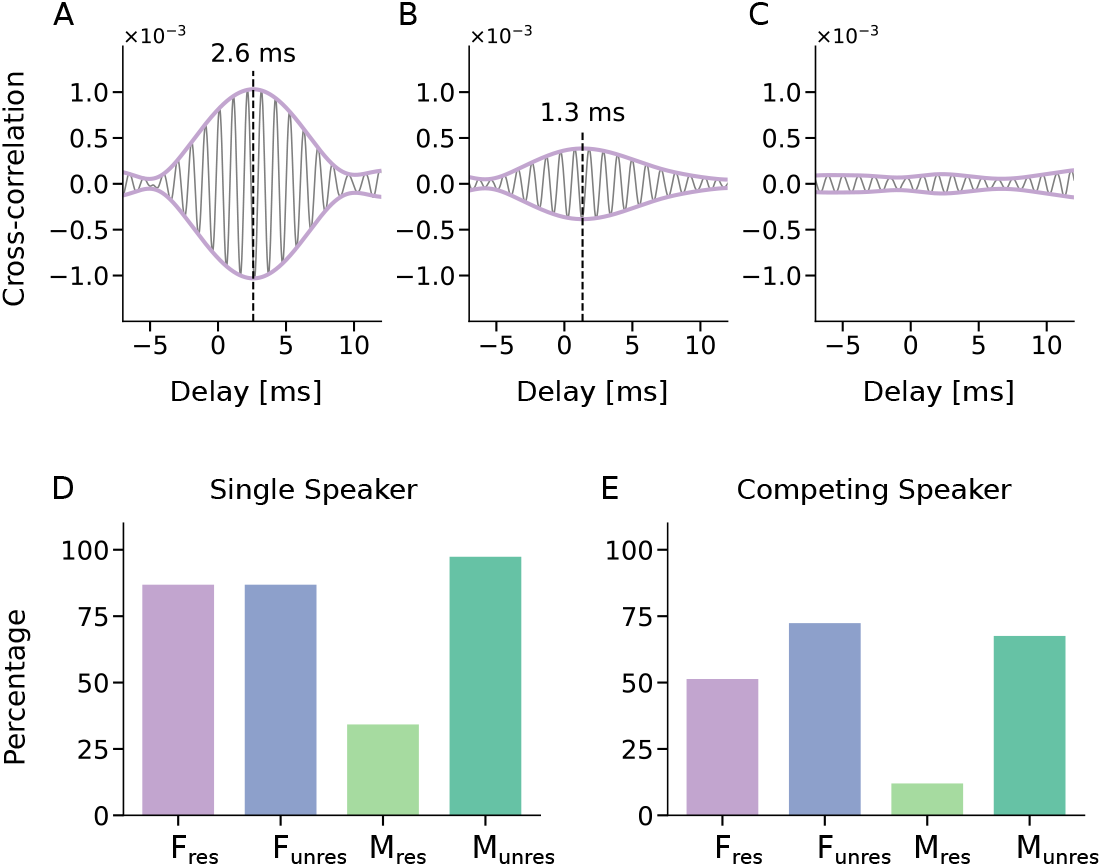
Recording of speech-like DPOAEs. Speech-like DPOAEs were measured through cross-correlating the expected DPOAE waveform *w*_2*n*−*m*_(*t*) with the microphone signal. (A,B) The envelope (purple) of the complex cross-correlation (black) exhibited a peak at a short delay (dashed line) that showed the presence of the speech-like DPOAE at this delay. (C) In contrast, if no peak emerged, the speech-like DPOAE could not be measured. The exemplary recordings show results from a single two-minute trial in the single-speaker scenario, for the *F*_res_ stimulus and for different subjects. (D,E), speech-like DPOAEs could be detected in most trials both in the single-speaker and in the competing-speaker scenario, except for the stimulus *M*_res_.

To assess how well the speech-like DPOAEs for the different stimuli could be measured, we computed the percentage of trials per stimulus type that yielded significant peaks in the cross-correlation. In the single-speaker scenario, the three stimuli *F*_res_, *F*_unres_, and *M*_unres_ all yielded significant speech-like DPOAEs in over 85% of single-speaker trials (Fig. 2D). However, the stimulus *M*_res_ performed notably worse, with only 35% of trials showing a significant speech-like DPOAE.

A similar pattern emerged in the competing-speaker scenario (Fig. 2E): the stimuli *F*_unres_ and *M*_unres_ performed well with speech-like DPOAEs detectable in about 70% of the trials, the stimulus *F*_res_ slightly lower, with about 50% of trials yielding significant speech-like DPOAEs, and *M*_res_ remained poor with only about 12% of trials producing a significant measurement.

To further compare the speech-like DPOAEs evoked by the different stimuli, we averaged the envelopes of the complex cross-correlations across trials and subjects, yielding grand averages.

In the single-speaker scenario, all stimulus types produced significant peaks, but with considerable variation in the amplitudes, that is, the values of the envelopes of the cross-correlations at the peak (Fig. 3E). The speech-like DPOAE for the stimulus *M*_res_, with an average frequency of 350 Hz, had the smallest amplitude. The highest amplitudes emerged for the stimuli *F*_res_ and *M*_unres_, both of which produced speech-like DPOAEs around 1 kHz.

**Figure 3.**
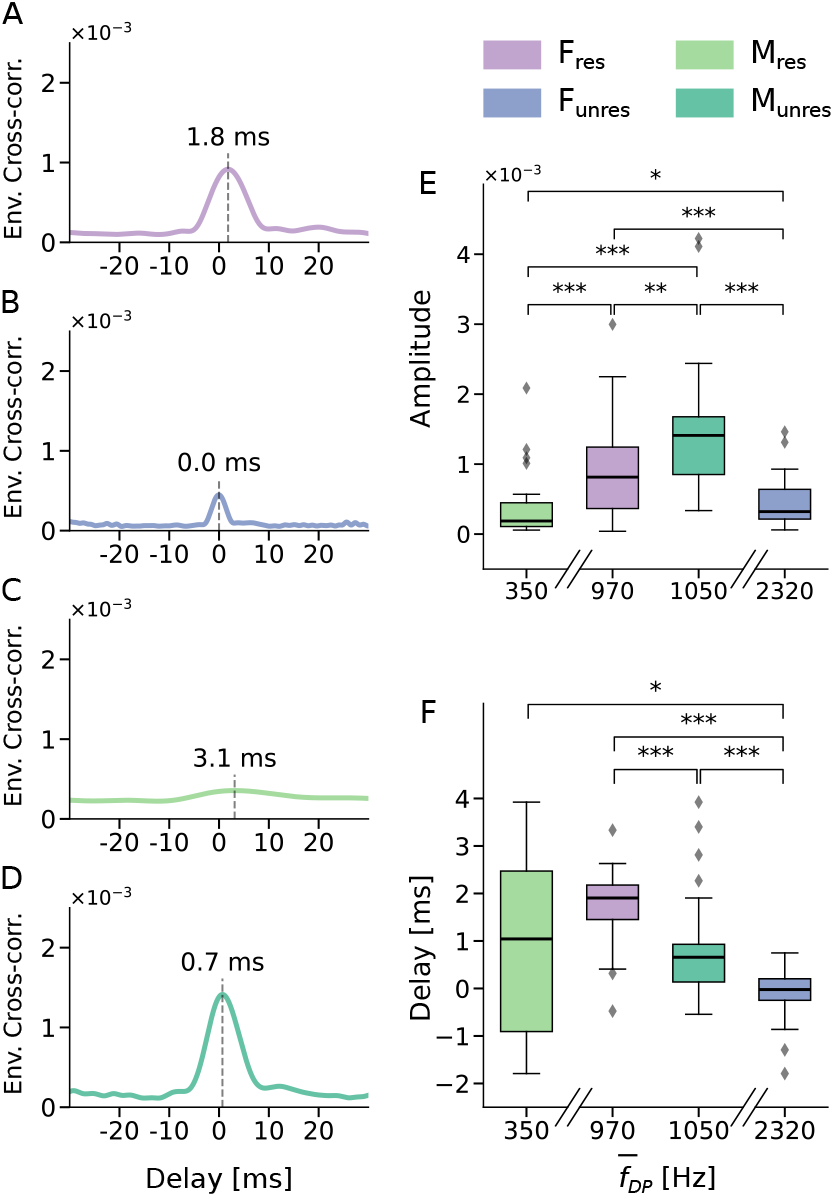
Speech-like DPOAEs in the single-speaker scenario. (A) – (D) Grand averages of the envelopes of the complex cross-correlations for the different stimuli. All peaks except the one for the *M*_res_ stimulus were clearly visible. The delays of the peaks are marked by dashed lines. E) The correlation coefficients at the peak latency varied significantly across stimulus types. They displayed an inverted U when plotted against the average frequency 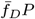 of the speech-like DPOAE, with the largest amplitudes emerging for speech-like DPOAEs around 1 kHz. F) Peak delays of individually identified peaks differed significantly across stimulus types. The variation in delays for the *M*_*res*_ stimulus was high due to the low number of significant peaks. Statistical significance is indicated as ^∗^ (*p* < 0.05), ^∗∗^ (*p* < 0.01), ^∗∗∗^(*p* < 0.001).

The delays of the speech-like DPOAEs also varied across the four stimulus types (Fig. 3F). The shortest delay was observed for the *F*_unres_ stimulus at 0 ms, while the longest delay occurred for the *M*_res_ stimulus at 3.1 ms. For the *M*_res_ stimulus, peak delays showed substantial variability across participants. This variability was likely due to the low number of significant peaks for this stimulus type (Fig. 2D–E), further underscoring the limited interpretability of the corresponding results.

Due to the small amplitude and the low rate of significant peaks observed, the stimulus *M*_res_ was excluded from further analysis.

For competing-speaker trials, the remaining stimuli (*F*_res_, *F*_unres_, *M*_unres_) produced significant peaks in all three attentional conditions (Fig. 4A-I). While the delays of the peaks remained stable across attentional conditions, they continued to vary between stimulus types.

**Figure 4.**
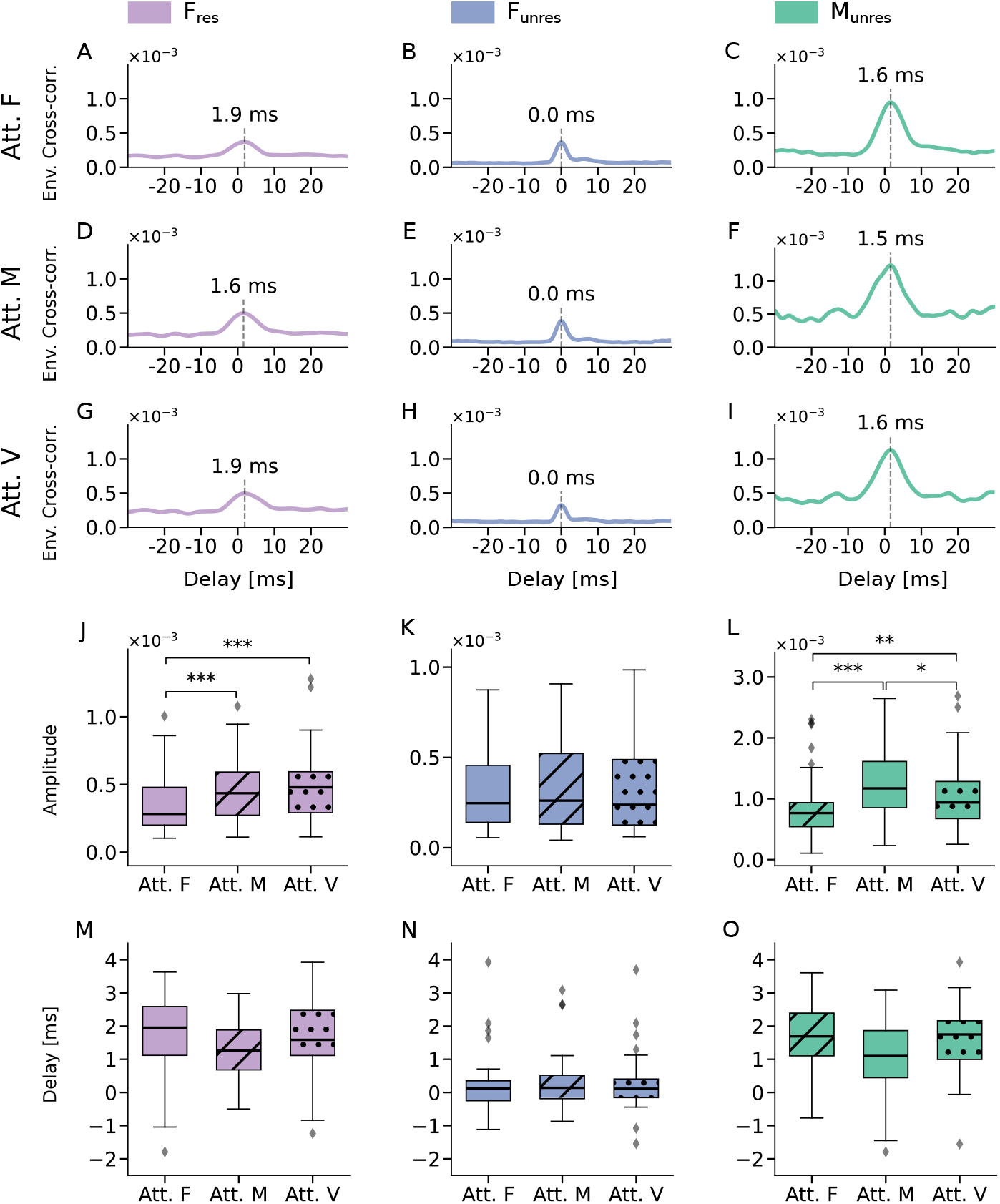
Attentional modulation of speech-like DPOAEs. (A) - (I) Envelopes of the complex cross-correlations of speech-like DPOAEs evoked by the three different stimuli *F*_res_, *F*_unres_, and *M*_unres_, in the three different attentional conditions attended female voice (Att. F), attended male voice (Att. M), and attended visual (Att. V.). All showed clear peaks at short delays (dashed lines). (J) - (L) Comparison of the peak amplitudes yielded attentional effects for the *F*_res_ stimulus and the *M*_unres_ stimulus, but not for the stimulus *F*_unres_. (M) - (O) The peak delays were not affected by the attentional focus, for neither of the three stimulus types. Statistical significance is indicated as ^∗^ (*p* < 0.05), ^∗∗^ (*p* < 0.01), ^∗∗∗^ (*p* < 0.001).

Out-of-ear control measurements did not yield significant peaks in the grand average for any stimulus type in single-or competing-speaker trials, confirming the absence of measurable distortion products when the probe was placed outside the ear canal.

### 3.3 Attentional modulation of speech-like DPOAEs

To assess effects of attentional focus on the speech-like DPOAEs in the competing-speaker scenario, we characterized the latter through both the delay of the peak in the complex cross-correlation and the amplitude, that is, the value of the envelope of the complex cross-correlation at the grand average peak delay.

We first quantified the influence of attention on the amplitude of the emissions. We started with the stimulus *F*_res_, that is, by assessing the speech-like DPOAEs evoked by resolved harmonics of the female voice (Fig. 4J). We found a significantly lower amplitude when the female speaker was attended (Att. F) than when the male speaker was attended (Att. M). The difference was highly statistically significant, with a *p*-value below 0.001 (*p* = 0.0003). The ratio of the amplitudes in the two conditions, Att. F. versus Att. M., was 0.8, or −2.2 dB.

The amplitude when attending the female speaker was also significantly lower than when reading the text, that is, when attention was focused on the visual modality, with a *p*-value below 0.001 (*p* = 0.0003, Fig. 4J). In this case, the amplitude ratio was 0.7, or −2.6 dB. In contrast, no significant difference emerged when comparing the attended male voice (Att. M) condition to the attend visual condition (Att. V).

For the speech-like DPOAEs elicited by unresolved harmonics of the female voice, the stimulus *F*_unres_, we did not observe any difference in amplitudes across the three attentional conditions (Fig. 4K).

For the male speaker, because the stimulus *M*_res_ did not give reliable speech-like DPOAEs, we could only assess the stimulus *M*_unres_ which utilized unresolved harmonics of the male voice. Significant amplitude differences emerged between all three conditions (Fig. 4L). The amplitude when attending the male voice was significantly higher than in the other two conditions. The ratio between the amplitudes when attending the male voice and when attending the female one was 1.5, or 3.8 dB (*p* = 7×10^−7^). In addition, the amplitude when ignoring the male voice, i.e. attending the female voice, was smaller than when reading the text (ratio of 0.8, or −2 dB; *p* = 0.002). The amplitude difference between attending the male voice and reading the text was slightly less pronounced (*p* = 0.03) with a ratio of 1.2, or 1.7 dB.

For the delays of the speech-like DPOAEs, we did not find any significant differences between the three attentional conditions, in neither of the three stimuli (Fig. 4M-O).

## 4 DISCUSSION

This study examined the feasibility of simultaneously eliciting and measuring multiple speech-like DPOAEs in response to four pairs of stimuli composed of resolved and unresolved harmonics that were derived from two distinct voices. We further investigated whether the speech-like DPOAEs were modulated by selective auditory attention as well as by intermodal attention, and whether such attentional modulation differed between resolved and unresolved harmonics.

Our study shows that four speech-like DPOAEs in response to multiple stimuli pairs can be measured successfully. However, we observed considerable variability in the quality of the recorded responses between the different speech-like DPOAEs (cf. Fig. 2), which may reflect either inherent variability in the generation mechanisms of these otoacoustic emissions or differences in noise levels due to varying frequencies of the eliciting stimulus waveforms. In addition, not all significant speech-like DPOAE peaks shared the same morphology; variations were observed in peak width, SNR, peak height, and peak latency.

While a previous study from our group already demonstrated the feasibility of recording speech-like DPOAEs [37], here, we extend this approach by eliciting speech-like DPOAEs with four simultaneously presented harmonic pairs instead of only one, thereby increasing both the speech-related information in the stimulus and the resulting DPOAE signal, and more closely approximating natural speech.

Further, we selected the employed harmonics such that they were either clearly resolved or clearly unresolved, strengthening the interpretability of our conditions. Indeed, our previous work reported indications of attentional effects in speech-like DPOAEs, but also noted inconsistencies between the male and female voices [37]. These may be attributed to differences in the harmonic structures employed, with the harmonics of the male voice in the previous study occupying the transition zone between resolved and unresolved harmonics.

Importantly, the auditory and visual stimuli remained statistically the same throughout the different attentional conditions. In particular, both audio streams and the visual text were presented concurrently with all four stimulus pairs for eliciting the speech-like DPOAEs. As a result, the only variable that differed between conditions was the participant’s attentional focus. This design minimized the likelihood of systematic confounds such as differences in movement or task structure. Furthermore, even if the middle-ear muscle reflex should have been activated despite the low stimulus volumes, it would have affected all attentional conditions equally.

Regarding the behavioral results, we found that the comprehension scores did not differ significantly across the attentional conditions, confirming that task difficulty was well-matched. Small differences in self-reported mental effort were likely attributable to speaker-specific factors such as intonation.

### 4.1 Frequency-dependency of speech-DPOAEs

Our evaluation of the amplitude of the speech-like DPOAEs in the single-speaker scenario revealed a pronounced dependency on the emission frequency (Fig. 3E). Consistent with previous findings on pure-tone DPOAEs, amplitudes were strongest for stimulus frequencies around 1 kHz [46]. Both lower and higher emission frequencies resulted in lower amplitudes. Because noise in electronics as well as in mechanical and acoustic systems increases at lower frequencies, the signal at the lowest emission, around 350 Hz for the stimulus *M*_res_, could not be detected in most trials. This stimulus was therefore excluded from further analysis.

### 4.2 Attentional Effects

We observed attentional modulation of the amplitudes of the speech-like DPOAEs for the stimuli *F*_res_ and *M*_unres_, but not for *F*_unres_.

The hypothesized differences between resolved and unresolved harmonics emerged clearly for the speech-like DPOAEs related to the female voice. Resolved harmonics, such as those employed for the stimulus *F*_res_, produce spatially distinct excitation peaks along the basilar membrane [34, 33]. Vibrations at these locations can be selectively enhanced through higher gain of the active process, which can help to enhance the neural representation of a target speech. In contrast, unresolved harmonics generate overlapping peaks along the basilar membrane, making such a spatial filter unfeasible. These considerations likely explain the presence of attentional modulation for the stimulus *F*_res_ together with its absence for the stimulus *F*_unres_.

The direction of the observed attentional effect for speech-like DPOAEs elicited by the stimulus *F*_res_, however, was unexpected: the resolved harmonics of the female voice counterintuitively caused *lower* amplitudes when attention was directed at the female voice versus when the male voice or the visual task was attended. Taken at face value, this result means that speech-like DPOAEs are weaker when a signal is attended than when it is ignored. This could suggest that the cochlea amplifies the harmonic structure of the target voice *less* than the background noise. Although unexpected, similar decreases have been reported in previous DPOAE studies [27, 47, 48] and occasionally elsewhere in the auditory system (e.g., decreases in neural tracking when intelligibility is high [49]).

Other studies also reported heterogeneous outcomes. Wittekindt et al. [30] found reduced DPOAE levels during visual attention when comparing levels to a baseline value of inattention, but no change during auditory attention, whereas Walsh et al. [26] observed attentional differences whose directions varied from subject to subject. These findings were challenged by Francis et al. [29], who observed that switching between states of attention and inattention produced changes in ear-canal noise that could mimic modulation of otoacoustic emissions. Our design avoids this confound: both auditory streams and the visual text were presented in all three attentional conditions, and only the focus of attention varied. We did not measure states of inattention. Thus, ear-canal noise and participant movement were expected to be equal across trials.

Other findings add to the mixed picture. Beim et al. [28, 50] reported higher SFOAEs during auditory attention in one study but failed to replicate the effect in a second cohort. Earlier work by Michie et al. [25] also found no attentional effect on tone-pip evoked OAEs. On the other hand, evidence for efferent modulation of otoacoustic emissions comes from a recent work showing that the predictability of tone sequences modulates DPOAE amplitudes depending on behavioral relevance [51]. Differences across study designs, types of OAEs, and susceptibility to noise likely contribute to these discrepancies.

The study most comparable to this is our earlier one on speech-like DPOAEs [37]. It found a positive attentional modulation coefficient for the female voice, indicating larger DPOAEs when the corresponding voice was attended. However, the reported effect was modest (*p* = 0.02) compared with the clearer differences observed here (*p* = 0.0003), and the actual DPOAE amplitudes did not differ significantly. Key methodological differences to this study include the previous use of only one harmonic pair per voice, partial placement of male harmonics in the resolved–unresolved transition region, and separate measurement blocks for attended and ignored states.

A possible origin of the weaker speech-like DPOAE when attending the corresponding voice may lie in the peculiar mechanics of the cochlea at low frequencies. Classical descriptions based on critical-layer absorption accurately capture basal, high-frequency processing [52, 18, 19], but several studies indicate that this framework may not apply straightforwardly at frequencies lower than 4 kHz [53, 54, 55, 56].

In this low-frequency regime, previous studies have proposed an alternative mode of operation, including independent resonance of the active process and unidirectional coupling between the basilar membrane and outer hair cells [57, 58]. Such mechanisms could suppress backward-propagating distortion products and may therefore provide an explanation for the direction of the attentional effects observed in our data.

The speech-like DPOAEs evoked by the *F*_res_ stimuli did not differ between the Att. M condition (equivalent to ignoring the female voice) and the Att. V condition. This suggests that cochlea activity at the resolved harmonics of an ignored speaker – in this case, the female voice – remains the same in these two conditions. One possibility, in line with the above considerations regarding apical cochlear mechanics, is that the resolved harmonics of a target speaker are enhanced through the active process, but yield lower speech-like DPOAEs, e.g. due to the peculiarities of low-frequency cochlear mechanics. When this voice is not attended, the harmonics are less enhanced, independent of whether attention is directed towards another auditory stream or the visual signal.

Because unresolved harmonics do not produce distinct peaks along the basilar membrane, the attentional modulation observed for the *M*_unres_ stimulus was unexpected. However, it is important to note that the *M*_unres_ stimulus was presented against a background of resolved harmonics of the female voice (Fig. 5). We hypothesize that attention to the female voice enhances cochlear activity in the regions corresponding to its resolved harmonics (purple shading in Fig. 5). This enhanced activity will also affect the responses to certain unresolved harmonics of the male voice, namely those that peak in the same section of the basilar membrane. These included the waveforms used in the *M*_unres_ stimulus. When the attentional focus changed from the female to the male voice, we hypothesize that cochlear activity in these regions became smaller, again affecting the corresponding unresolved harmonics of the male voice, including the *M*_unres_ stimulus. The attentional modulation for the *M*_unres_ stimulus was thus likely caused by changes in cochlear activity for the resolved harmonics of the female voice.

**Figure 5.**
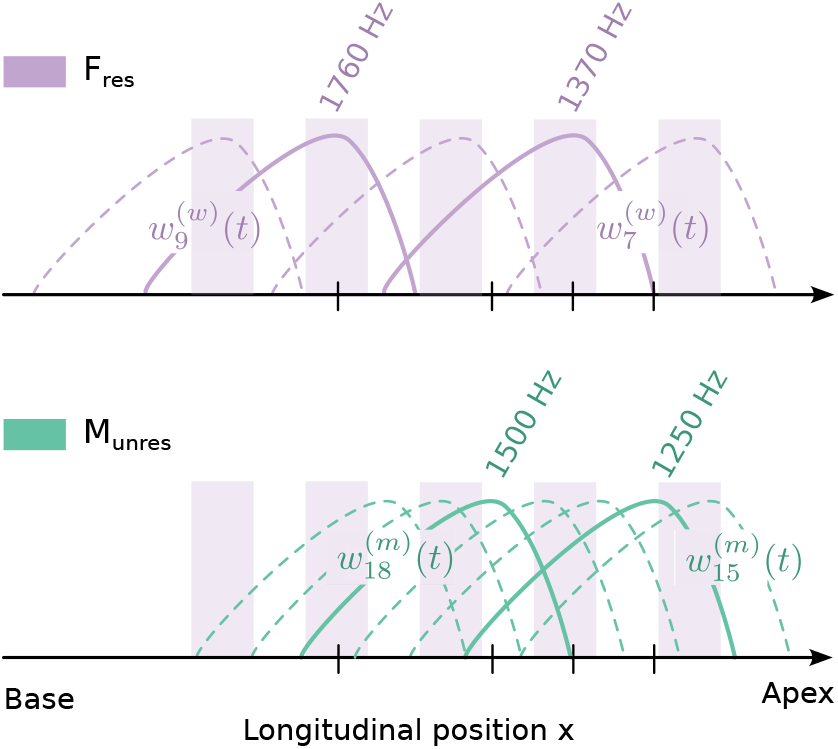
Resolved and unresolved harmonics. In the part of the cochlea where the *F*_res_ and the *M*_unres_ stimuli have their characteristic places, the harmonics of the female voice are resolved (top) while those of the male voice are unresolved (bottom). The cochlear active process may enhance the response at the locations where the resolved harmonics of the female voice peak (purple shading). This enhancement will then also affect the responses to the unresolved harmonics of the male voice at these locations.

### 4.3 Delay of the speech-DPOAEs

As expected, the delays of the speech-like DPOAEs did not depend on the focus of selective attention (Fig. 4M-O). However, the delays differed between the different stimulus types as assessed in the single-speaker scenario. Speech-like DPOAEs evoked by the *F*_res_ stimulus peaked around 2 ms, corresponding to approximately 2 cycles on the basilar membrane. Because the traveling wave, for low frequencies, accumulates about one cycle between the base and the peak location, this delay matches with the expectation of the stimuli propagating towards the peak location, the active process generating distortion there, and the latter propagating back through a basilar-membrane wave [59, 60, 46, 61, 18].

In contrast, the speech-like DPOAEs evoked by the *F*_unres_ and the *M*_unres_ stimuli occurred at shorter latencies, 0 ms respectively 0.7 ms, corresponding to approximately 0 and 0.7 cycles. This suggests that these distortions are created basal to the peak of the corresponding basilar-membrane waves, and that they might propagate back from their generation site through a fast wave such as a compression wave [62, 63, 64].

The *M*_*res*_ stimulus did not give reliable responses and was hence excluded from more detailed analysis. It only produced a low percentage of significant peaks (Fig. 2D,E), weakly pronounced grand average peaks (Fig. 2D-E and Fig. 3C), as well as large variation in delays (Fig. 3F).

#### Limitations

Comparability of the stimuli derived from the male voice was limited. In the competing speaker scenario, the stimulus *M*_res_ failed, in most trials, to produce significant speech-like DPOAEs for statistical evaluation. Additionally, the stimulus *M*_unres_ overlapped in frequency with the stimulus *F*_res_, compromising interpretability due to potential confounds in the origin of observed effects. These limitations suggest that, for the speech-like DPOAEs, the average stimulus frequencies should exceed 1 kHz to ensure reliable measurements. Moreover, when aiming at simultaneous stimulation with different harmonic combinations and multiple speakers, avoiding frequency overlap between stimuli is desirable to support a clear interpretation of the results.

### 4.4 Conclusion

Our study shows that attention modulates the morphology of speech-like DPOAEs elicited by multiple, simultaneously presented harmonic pairs derived from natural speech signals. These findings indicate that DPOAEs likely also arise in response to natural, running speech, even though extracting such responses remains challenging due to the high noise floor and spectral complexity of real speech. Our experimental paradigm offers nonetheless a promising approximation for probing cochlear responses to ecologically valid speech stimuli.

We observed attentional effects for both resolved and unresolved harmonics. However, we argue that the effects seen for the unresolved harmonics most likely resulted from attentional modulation of the resolved harmonics of the competing speaker. Importantly, the direction and pattern of attentional modulation were consistent, whether attention was shifted from the target speech to a competing voice or from the target to a visual task.

Unexpectedly, attention reduced – rather than enhanced – the speech-like DPOAE associated with the attended speaker. Given previous heterogeneous findings in the literature, we aim to replicate this effect in an upcoming study to establish the robustness and generalizability of our results.

More broadly, we hope that the speech-like DPOAE paradigm introduced here will contribute to a deeper understanding of how the active cochlea participates in the perceptual and cognitive processing of complex naturalistic signals such as speech.

## DATA AVAILABILITY

The datasets generated and analysed during the current study are available in the FAU speech-like DPOAE repository on the platform zenodo, https://zenodo.org/records/16837673.

## CODE AVAILABILITY

All custom code used for stimulus generation, speech-like DPOAE analysis via cross-correlation, and statistical evaluation is openly available on GitHub at https://github.com/janna-stb/dpoae_attention_study.git under the MIT License.

## FUNDING

This work was funded by the Deutsche Forschungsgemeinschaft (German Research Foundation) through grant 514955521 (to T.R.).

## AUTHOR CONTRIBUTIONS

J.S. and T.R. designed research and analyzed data; J.S. performed research and wrote the paper; T.R. revised the paper.

## AUTHOR COMPETING INTERESTS

The authors declare no competing interests.

## Notes

https://zenodo.org/records/16837673

https://github.com/janna-stb/dpoae_attention_study.git

### Competing Interest Statement

The authors have declared no competing interest.

### Summary of Updates

We partly revised the manuscript to improve clarity and contextualization. First, several methodological descriptions were expanded or clarified, particularly regarding the generation of speech-derived stimuli, the construction of harmonic waveforms, and the computation of speech-like DPOAEs. These revisions aim to enhance reproducibility and ensure that the signal-processing steps are transparent to readers. Second, the Discussion section was restructured and refined. We strengthened links to existing literature, added missing comparisons with previous DPOAE attention studies, and streamlined sections that were previously considered too speculative. Finally, minor stylistic and organizational edits were made across the manuscript to improve coherence and narrative flow. Importantly, no analyses or results were changed relative to the original version; all revisions are solely clarifications and improvements in presentation.

